# Evaluation of 20 enset (*Ensete ventricosum*) landraces for response to *Xanthomonas vasicola* pv. *musacearum* infection

**DOI:** 10.1101/736793

**Authors:** Sadik Muzemil, Alemayehu Chala, Bezuayehu Tesfaye, David J. Studholme, Murray Grant, Zerihun Yemataw, Temesgen Magule Olango

## Abstract

Bacterial wilt, caused by *Xanthomonas vasicola* pv. *musacearum* (Xvm), formerly *X. campestris* pv. *musacearum*, is the most threatening and economically important disease of enset (*Ensete ventricosum*), the multipurpose food security crop orphan to south and southwestern Ethiopia. Xvm has also had a major impact on banana and plantain production in East Africa following its detection in Uganda in 2001 and subsequent spread. Effective control of this disease currently relies on integrated disease management (IDM) strategies including minimization of field pathogen inoculum and deployment of wilt resistant enset landraces. Identifying landraces with stable and durable Xvm resistance will greatly accelerate breeding of varieties that can be included as a component of IDM. In this study, 20 enset landraces previously reported to exhibit lower susceptibility to Xvm were grown in pots under open field conditions and inoculated with an aggressive Xvm inoculum isolated from a disease hotspot area. Longitudinal and survival analyses were applied to each landrace, based on disease units representing a combination of area-under-disease progress stairs, disease index and apparent infection rate. Considerable variation was observed among the 20 landraces; however, none exhibited full immunity to Xvm infection. Three landraces, viz. Hae’la, Mazia and Lemat (HML), showed lowest susceptibility to Xvm as evidenced by lower disease units and higher survival rates. Landraces Kuro, Gezewet, Bededet, and Alagena showed similar levels of Xvm infection as did HML, but with lower survival rates. By contrast, landrace Arkia showed the highest infection level and lowest survival rate, suggesting a high degree of susceptibility to Xvm. This study identifies new material that can be used in future breeding programmes to develop Xvm-resistant enset varieties.

## 1. Introduction

Enset (*Ensete ventricosum* (Welw.) Cheesman) is a diploid (2n=18), herbaceous, perennial monocarpic crop belonging to the family *Musaceae*. Enset is often referred to as false banana, due to its phenotypic resemblance to banana (*Musa* species). The crop is cultivated exclusively in Ethiopia, where it has a considerable economic and social importance for millions (Brandt *et al.*, 1997; Borrel et al., 2019).

Enset is tolerant to prolonged drought and provides a year-round source of staple nutritious food. It is thus widely cultivated across south and southwestern Ethiopia contributing to improved food security for more than 20 million Ethiopians (Yesuf and Hunduma, 2012, Yemataw et al., 2017). For farmers, enset is more than a year-round staple food, as it provides multiple additional daily benefits yet requires little crop management husbandry. The multipurpose benefits are derived from different enset landraces that are particularly suited for feed, fiber, packaging, construction material as well as providing a medicinal role (Brandt et al., 1997; Nurfeta et al., 2008; Yemataw et al., 2017). Due to its long history of cultivation across diverse ethnic groups, enset has significant cultural and socio-economic value in Ethiopia (Shigeta, 1990; Olango et al., 2014, Borrell et al., 2019).

Enset production is threatened by bacterial wilt disease caused by *Xanthomonas vasicola* pv. *musacearum* (Xvm) (Ashagari, 1981, 1985; Archido and Tessera, 1993; Tessera et al., 2008; Yesuf and Hunduma, 2012; Nakato et al., 2018). The bacterial disease was first reported in Kaffa district of Ethiopia in 1960’s initially on enset (Yirgou and Bradbury, 1968) and subsequently on banana (Yirgou and Bradbury, 1974). However, the first observations for the bacterial disease on enset in Ethiopia dates back to 1930s (Castellani, 1939). The causative agent was previously known as *Xanthomonas musacearum* and *X. campestris* pv. *musacearum*, but a recent taxonomic study been transferred it to the species *Xanthomonas vasicola* (Studholme et al. 2019 in press). Currently, the disease is found distributed in all enset growing areas of southern and southwestern Ethiopia, where it has a devastating impact on enset production (SARI-McKnight CCRP, 2013; Blomme et al., 2017; and Nakato et al., 2018). Other diseases caused by fungi (Tessera and Quimio, 1993), viruses (Tessera et al., 2003) and nematodes (Bogale et al., 2004) also affect enset. Mammals and pests such as porcupines, mole rats, wild pigs and insects such as mealybugs (Azerefegne et al., 2009) also impact enset production (Handoro et al., 2012).

Enset landraces in Ethiopia might offer enhanced resistance to Xvm, thus potentially offering a huge genetic resource to farmers though no systematic patho-testing for responses to Xvm has yet been implemented. Currently, farmers claim that certain enset landraces show relatively low level of infection to Xvm and they often incorporate these landraces in their backyard landrace mixture as one option for disease management (Ashagari, 1985; SARI-McKnight CCRP, 2013; Yemataw et al., 2016). Nevertheless, identification of resistant landraces has been challenging. Literature has reported enset landraces with lower but varying susceptibility to Xvm infection (Ashagari, 1985; Archaido and Tessera, 1993; Handoro and Welde-Michael, 2007; Welde-Michael et al., 2008; Haile et al., 2014; Hunduma, 2015; Wolde *et al.*, 2016; Handoro and Said, 2016). However, inconsistency in enset landraces’ responses to Xvm has been observed (Ashagari, 1985; Handoro and Welde-Michael, 2007; Welde-Michael *et al.*, 2008; Tadesse *et al.*, 2008), with variation in virulence among batches of Xvm inoculum being the most probable cause. Notably, existing Xvm resistance phenotyping data primarily comprise assessments of individual landraces.

In banana transgenic approaches have been demonstrated for resistance against Xvm (Namukwaya et al., 2012) due to absence of genetic resistance in the cultivated banana germplasm pool. While initial results look promising, this approach would require biosafety regulations for the adoption and use of transgenic plants, and adoption of the approach to enset. Hence, the availability of sources of resistance to Xvm in enset germplasm could serve as a breeding stock against Xvm in enset as well as banana plant.

To date, no systematic screening has been undertaken on the same enset landraces that are reported to have high tolerance to Xvm infection. Such a study would greatly assist breeding/selection efforts to identify elite landraces exhibiting enhanced Xvm resistance. In this paper we have undertaken field experiments to re-evaluate selected enset landraces previously reported to have reduced susceptibility to Xvm. We undertook a detailed study of the infection reaction and identified specific enset landraces with potential to contribute towards sustainable management of the disease, using a common garden experiment at a single site to minimize environmental variability and enable a detailed study of the infection reaction and identify specific enset landraces with potential to contribute towards sustainable management of the disease.

## 2. Materials and Methods

### 2.1. Description of the study site

Evaluation of enset landraces for resistance to *Xanthomonas vasicola* pv. *musacearum* (Xvm) was carried out using an open-field potted experiment at Southern Agricultural Research Institute (SARI), Hawassa, Ethiopia. The site is located at 7°4’ N and 38°31’ E with an elevation of 1700 meters above sea level and having an annual rain fall of 1100 mm. The annual average, minimum, and maximum temperatures at Hawassa are 20.6°C, 13.5°C and 27.6°C, respectively.

### 2.2. Plant material acquisition and multiplication

This study used 18 enset landraces previously reported to exhibit low level infection or having contrasting infection phenotypes following Xvm inoculation. Landrace ‘Arkia’, was also included for comparison because it was previously reported to support a high level of Xvm growth. Also included was landrace ‘Bota Arkia’, which has a similar name to landrace ‘Arkia’ but of different origin. This gave a total 20 landraces for the present study (Table 1).

**Table 1.** Enset landraces studied for their reaction to *Xanthomonas vasicola* pv. *Musacearum* (Xvm)and previously recorded reaction to Xvm.

| No | Enset landraces | Collection zones* | Previously reported response** | References for landrace reaction to Xvm |
| --- | --- | --- | --- | --- |
| 1 | Abatemerza | Kambata | Resistance /Tolerant | Handoro and Said (2016) |
| 2 | Ado | Sidama | Moderately Tolerant | Ashagari (1985), Welde-Michael et al. (2008) |
| 3 | Alagena | Wolaita | Resistance /Tolerant | Handoro and Said (2016) |
| 4 | Arkia | Wolaita | Susceptible | Handoro and Said (2016) |
| 5 | Bededet | Gurage | Moderately Tolerant | Wolde et al. (2016) |
| 6 | Bota Arkia | Dawro | NA |  |
| 7 | Dirbo | Kambata | Resistance /Tolerant | Handoro and Said (2016) |
| 8 | Gefetano | Wolaita | Resistance /Tolerant | Handoro and Said (2016) |
| 9 | Genticha | Sidama | Moderately Tolerant | Ashagari (1985), Welde-Michael et al. (2008) |
| 10 | Gezewet | Gurage | Susceptible | Welde-Michael et al. (2008) |
|  |  | Gurage | Resistant | Wolde et al. (2016) |
| 11 | Ginbuwa | Gurage | Tolerant | Handoro and Said (2016) |
| 12 | Godere | Wolaita | Resistance /Tolerant | Handoro and Said (2016) |
| 13 | Hae'la | Kambata | Highly Tolerant | Tessera et al. (2000), Gizachew et al. (2008) |
| 14 | Kuro | Dawro | Moderately Tolerant | Handoro and Said (2016) |
| 15 | Lemat | Gurage | Moderately Resistance /Tolerant | Welde-Michael et al. (2008), Handoro and Said (2016) |
|  | Lemat | Gurage | Susceptible | Mekuria et al. (2016) |
| 16 | Mazia | AARC (Originally Collected from Dawro) | Resistance /Tolerant | Handoro and Welde-Michael, (2007), Welde-Michael et al. (2008), Tessera et al. (2008), Tariku et al. (2015), Handoro and Said (2016) |
| 17 | Nechuwe | Gurage | Moderately Tolerant | Welde-Michael et al. (2008), |
|  |  | Gurage | Susceptible | Wolde et al. (2016) |
| 18 | Unjeme | Kambata | Resistance /Tolerant | Handoro and Said (2016) |
| 19 | Wachiso | Kambata | Resistance /Tolerant | Handoro and Said (2016) |
| 20 | Yesha | Dawro | Resistance /Tolerant | Handoro and Said (2016) |

\* Administrative zone in Southern Ethiopia where the respective enset landrace was collected from farmers field. Original collection history of the landraces obtained from the articles listed.
\*\* The recorded Xvm response indicated was derived from the accompanying cited article(s).

To initiate the experiment, underground corms of 2-3 year old mother plants from each of the 20 landraces were macro-propagated to produce true-to-type suckers for the experiment. Mother corms of landrace Mazia were kindly provided by Areka Agricultural Research Center (AARC). The remaining 19 enset landraces were collected from enset farmers in six enset growing zones of southern Ethiopia viz. Dawro, Gedio, Gurage, Kambata, Sidama and Wolaita. Collection of the landraces from these areas was undertaken by first confirming that the landrace described was true for its reaction to Xvm, followed by a rapid appraisal on site to avoid the possiblilty of mis-identification (Figure 1).

**Figure 1.**
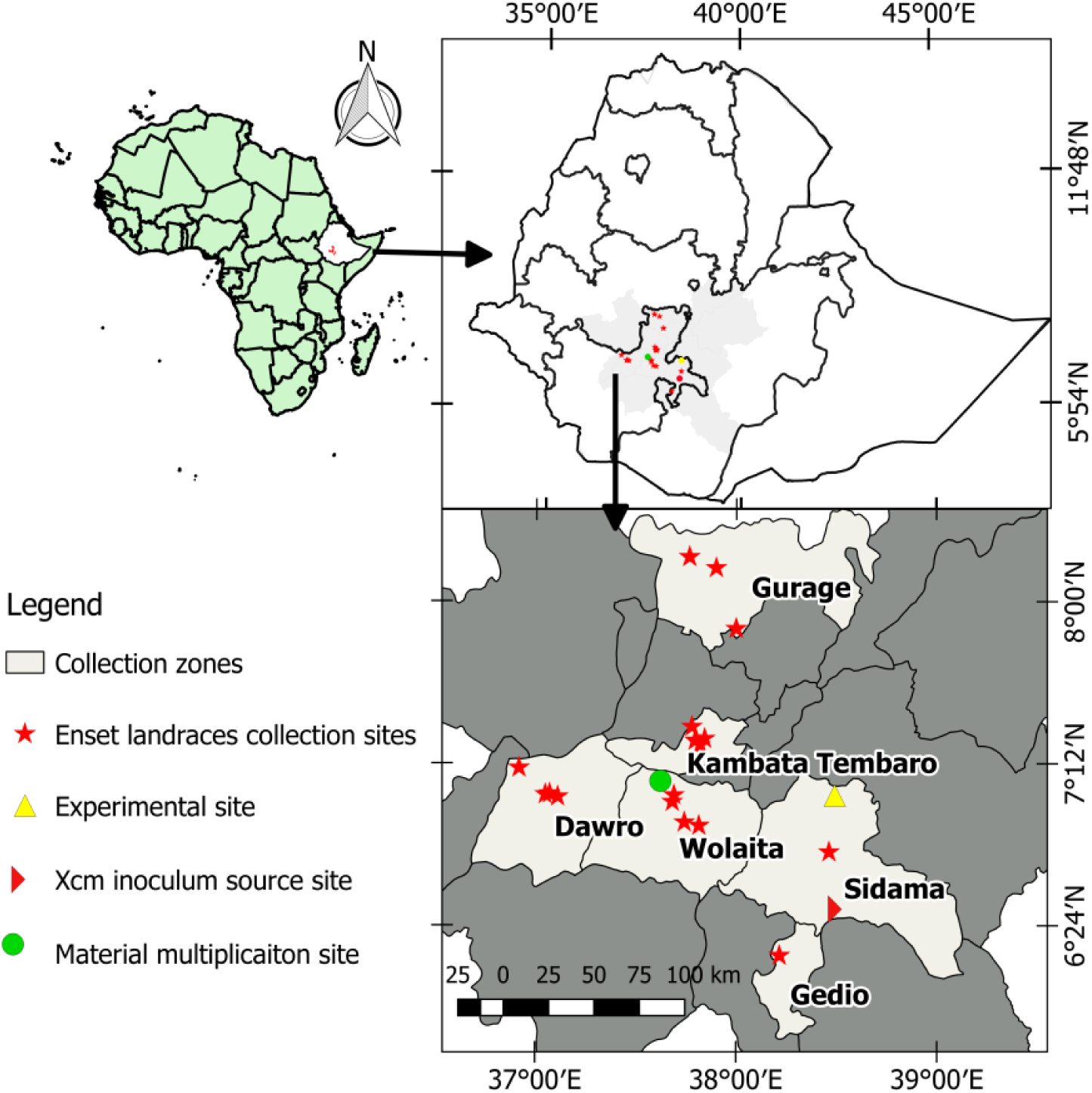
Geographical locations of enset landraces collection and experimental sites.

Following collection, enset landraces were multiplied at Areka Agricultural Research Center (AARC) using an enset macro-propagation method previously described (Yeshitila et al. 2009). Briefly, the mother corm was dissected into two equal halves and the apical tissue removed from the center of each half corm to allow secondary meristems to develop into suckers for planting. These dissected corms are dried in the shade for 3 hours prior to planting. Each half corm was planted in slanted orientation in holes of 1 m × 1m and covered with soil.

After 12 months, multiplied enset suckers were uprooted and transplanted into a 5 kg capacity plastic pot filled with sun dried mixtures of soil: sand: manure at a ratio of 3:1:1 (Quimio, 1992) and placed in an open field experimental site at SARI. Suckers were allowed to establish for two months to get sufficient leaves (>3) to allow inoculation with Xvm. During this establishment period, suckers in plastic pots were watered daily prior to inoculation with Xvm and for a subsequent two months after Xvm inoculation. Watering thereafter was reduced to two times per weeks for remaining experimental period.

### 2.3. Preparation of bacterial suspension for inoculation

Virulent Xvm was collected from a disease hot spot in the Hagereselam area, Sidama Zone, southern Ethiopia. Xvm bacterial ooze from young leaves and /or pseudostem of diseased enset plants was harvested into sterile distilled water and preserved at 4 °C until use. Two sets of bacterial suspensions were prepared for hypersensitivity and pathogenicity tests. One comprised the uncultured bacterial suspension. The other was derived from a day-old preserved field harvested Xvm isolates streaked on YPSA (5 g yeast extract, 10 g peptone, 20 g sucrose and 15 g agar) – a growth and isolation medium used for selecting pure Xvm colonies (Haile *et al.*, 2014). These plates were incubated at 28°C for 24 h (Schaad and Stall, 1998). Single colonies with a yellow, convex, mucoid morphology typical of Xvm were harvested and preserved in YPSA slants at 4 °C.

### 2.4. Hypersensitivity and pathogenicity tests

Both uncultured and cultured suspension were tested for hypersensitivity and pathogenicity on two-month-old tobacco (*Nicotiana tabacum*) or the highly susceptible control enset landrace ‘Arkia’. In hypersensitivity test conducted on *N. tabacum* (Bobosha, 2003) using both inoculum types independently, 2 ml of a bacterial suspension containing ∼1.0 × 10^8^colony forming units (cfu) per mL (approximately OD_600_ = 0.5) was used for inoculation. A positive hypersensitive response was scored if tissue exhibited yellow clearing chlorosis limited to around the point injection.

For initial assessment of pathogenicity tests, 14-month-old (i.e. two months after transplanting) disease-free enset suckers of the susceptible landrace ‘Arkia’ (Handoro and Welde-Michael, 2007) were infected with 4 mL of bacterial suspension (∼10^8^ cfu/mL at OD_600_ = 0.5) from uncultured and cultured inocula.

### 2.5. Inoculation of Xvm to test landraces

As the uncultured bacterial suspension resulted in shorter incubation period and more severe disease on both *N. tabacum* and enset landrace Arkia during the pathogenicity tests, field-harvested, uncultured Xvm suspensions preserved at 4 °C were used as inoculum for landrace evaluation. Suckers of 14-months-old enset landraces (two months after transplanting in potted soil mix) were inoculated with a 4 mL aliquot of the bacterial suspension, adjusted to ∼10^8^ cfu/mL as described above, by infiltration with a hypodermic sterile syringe into the youngest innermost leaf petiole (Figure 2). A new sterile hypodermic syringe was used for inoculating each sucker of every landrace. Control plants were infiltrated with the same volume of sterile distilled water. The pot experiment was organized in four replicates of 20 landraces, each landrace containing 10 plants and landraces within each replication were randomized. Thus, the entire experiment comprised a total of 800 plants. This included a total of 30 individuals of the 20 enset landraces inoculated with uncultured Xvm suspension and a further 10 individuals of the 20 plants comprising negative controls inoculated with sterile distilled water.

**Figure 2.**
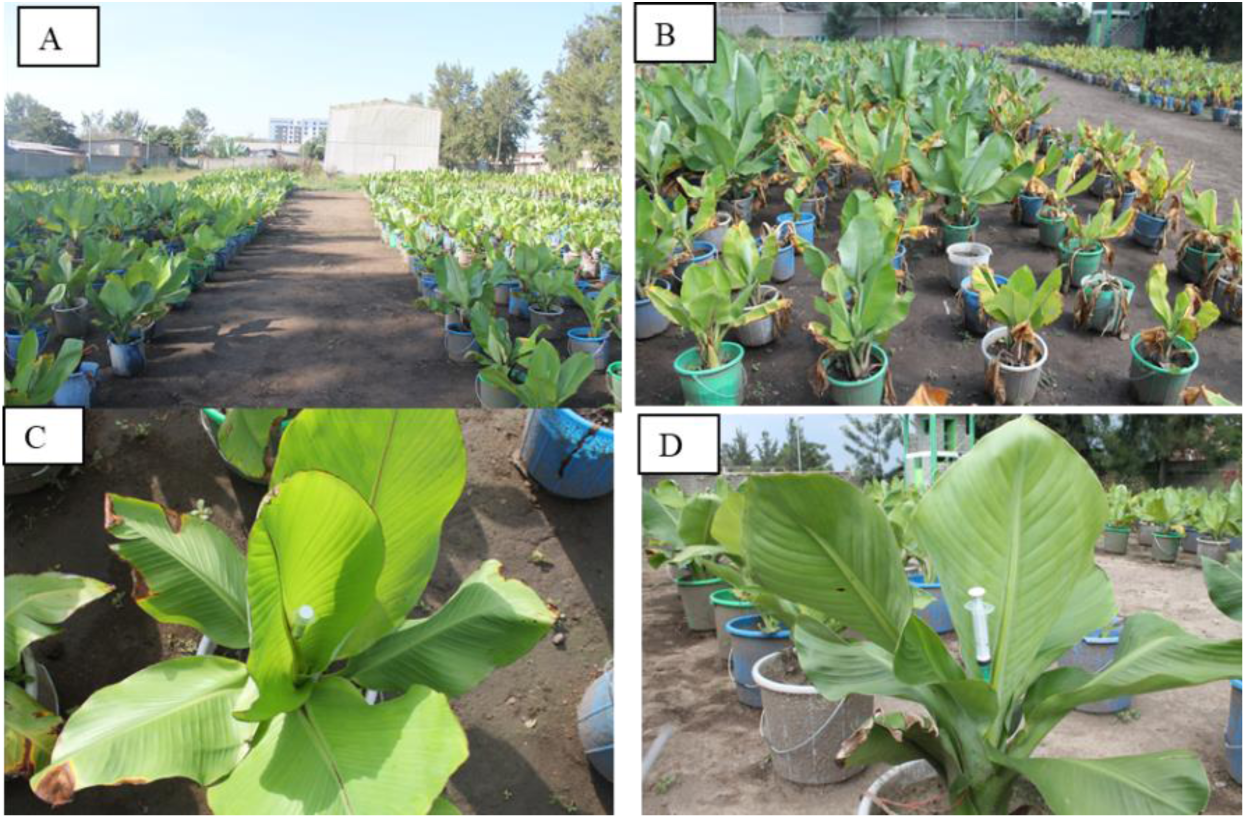
Evaluation of enset landraces grown under potted soil mix (A and B) and inoculation of landraces (C and D) with uncultured *Xanthomonas vasicola* pv. *musacearum* (Xvm) suspension

### 2.6. Measurements and data analysis

#### Data recording and quantifying disease progression

Data collection was initiated a week after inoculation and continued weekly for the first three weeks, then at two-week intervals for 155 days thereafter. It was notable that in the first two weeks following infection, some enset landraces showed unusual infection responses including twisting of the leaf blade in some landraces, rolling or curling of the leaf tip and leaf edge in other enset landraces. From the third week after Xvm infection onward, we observed typical wilting symptoms such as severe drooping of the top 25% of leaf blade, collapsing of the leaf blade, wilting of both inoculated and adjacent uninoculated leaves, chlorosis across the majority of leaves and, in certain landraces, complete death. At each evaluation time, the number of leaves per plant that showed these consistent wilting symptoms and total number of asymptomatic leaves per plant were recorded. Data from the “inconsistent” first two-week period were excluded from analysis as it ws not possible to associate these initial responses with a phytopathological response.

Four response variables were used to quantify enset disease phenotypes following Xvm infection: disease index (DI), area under disease progress stairs (AUDPS), apparent infection rate (AIR) and survival. DI is the percentage of symptomatic leaves per individual at each evaluation period averaged across the 4 replicates (Schandry, 2017). DI helps to quantify disease symptoms over the evaluation period using a scale of 0-4, with 1 and 4 corresponding to 25% and 100% of total wilted leaves per plant respectively.

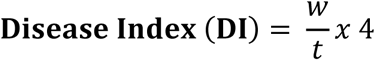

Where, *w* is the number of symptomatic leaves and *t* is the number of total leaves of a single plant. AUDPS (Simko and Piepho, 2012) was used to estimate disease accumulation and progress over time. AUDPS is proposed to provide better estimates of the disease by giving weight closer to the optimal than that derived from AUDPC (area under disease progress curve) assessments (Madden *et al.*, 2007). AIR, corresponding to the speed at which an epidemic develops (Meena *et al.*, 2011), was calculated as the slope of disease index development. Finally, survival analysis was applied to study the fraction of survivors per time point among enset landraces challenged with Xvm. Survival data was generated according to Schandry (2017) with a customized DI cut-off point of 2.23. This cut-off DI was used for comparison purpose and is the maximum DI value of landrace Mazia that is frequently cited (Table 1) for its resistance/tolerance to Xvm.

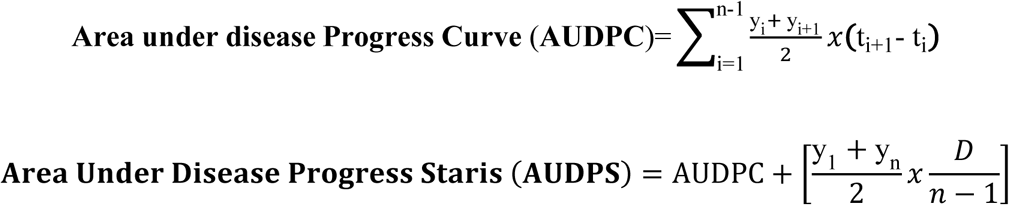

Where, y_i_ is disease index at the i^th^ observation, t_i_ is time in days at the i^th^ observation, n is the number of total observations, and D is duration from the first to the last observation (D=t_n_-t_1_)., n is the number of total observations, and D is duration from the first to the last observation (D=t_n_-t_1_).

#### Data analysis and visualization

Various functions from the following R packages were used for data manipulation and formatting: *broom* (Robinson et al, 2015), *dplyr* (Wickham and Francois, 2015), *magrittr* (Bache and Wickham, 2014), *modelr* (Wickham, 2016a), *stargazer* (Hlavac, 2015), *stringr* (Wickham, 2010), *tidyr* (Wickham et al., 2017) and *tidyverse* (Wickham, 2016b). For most of the disease parameters in this study, data were fitted to linear mixed model using functions in *lme4* package (Bates et al., 2015) to properly account for differential disease development in enset landraces over time. The linear model (lm) function from the base *stats* package was employed. For model comparison in the disease units and survival analysis, Akaike information criterion (AIC) and Schwarz’s Bayesian information criterion (BIC) were used to identify the best model among the alternatives. Models with the lower AIC and BIC value were used for analysis. Functions from *emmeans* (Russell, 2019) and *lmerTest* (Kuznetsova *et al.*, 2016) were used to extract means from data fitted to linear and linear mixed models respectively, applying the process “data manipulation for result visualization”. Graphic visualizations were done using *ggplot2* (Wickham, 2009).

Functions from the following packages were used for analysis: *MESS* (Ekstrøm, 2016), *survival* (Therneau and Lumley, 2011), *survcomp* (Schröder et al., 2011), *rms* (Harrell Jr, 2016), coxme (Therneau, 2015b), *lme4* (Bates et al., 2015), *lmerTest* (Kuznetsova *et al.*, 2016), *multcomp* (Hoth on *et al.*, 2008), and *rcompanion* (Mangiafico, 2017). Packages *rmarkdown* (Allaire et al., 2015) and *knitr* (Xie, 2014, 2015, 2016) were used to generate R Markdown (.rmd) files. Furthermore, scripts used in data manipulation, analysis and visualization are available in the Rmarkdown (**supplementary R_script and Enset_Xvm_data**) which contains a full description of data management and analysis.

Generalized linear hypothesis testing, adjusted for multiple comparisons using Tukey’s method (Hothorn et al., 2008), was used to assess statistically significant differences at *P*-value cut-off of 0.05. Statistical significance among treatments was determined using the “compact letter displays (cld)” method in the graphic visualization; cld function from *multcomp* (Hothorn et al., 2008) package. Treatments within the same “letter group” are not significantly different whereas treatments with different letters display a significant difference.

## 3. Results

### 3.1. Hypersensitivity and pathogenicity tests

A typical hypersensitive response (HR) was observed on tobacco leaves from both of uncultured and medium based Xvm suspension. However, uncultured Xvm suspension caused a severe HR on tobacco within three days post inoculation whereas symptoms were delayed for up to eight days when cultured Xvm was used (data not shown). The effect of inoculum type on symptom appearance was more pronounced during pathogenicity testing on the susceptible landrace Arkia. It took 21 days after inoculation (DAI) with uncultured Xvm suspension to cause wilting and complete collapse of enset leaves of the susceptible landrace Arkia whereas the same disease level was not attained in the cultured suspension inoculation until 30 to 45 DAI.

### 3.2. Symptom description

A range of symptoms was observed during the course of infection and subsequent disease development on Xvm-challenged enset landraces. Necrosis around the point of inoculation and surrounding tissues was observed 3 DAI in most landraces (Figure 3). At early stages of infection, up to 15 DAI, landraces showed a varying range of symptoms. Included among these symptoms were twisting and slight leaf curling, and drooping of the blade and tip of the inoculated leaf. The leaf blade around the Xvm inoculated area often became deformed, twisted or curving inwards. These symptoms were replaced by severe curling of the leaf edge, drooping and folding back of leaf blade from 15 DAI, symptoms that were consistently observed in all landraces. Gradually, drooping from the leaf apex and folding back or collapsing of leaves became the most prominent symptoms as the disease developed. All tested enset landraces showed one or more these symptoms.

**Figure 3.**
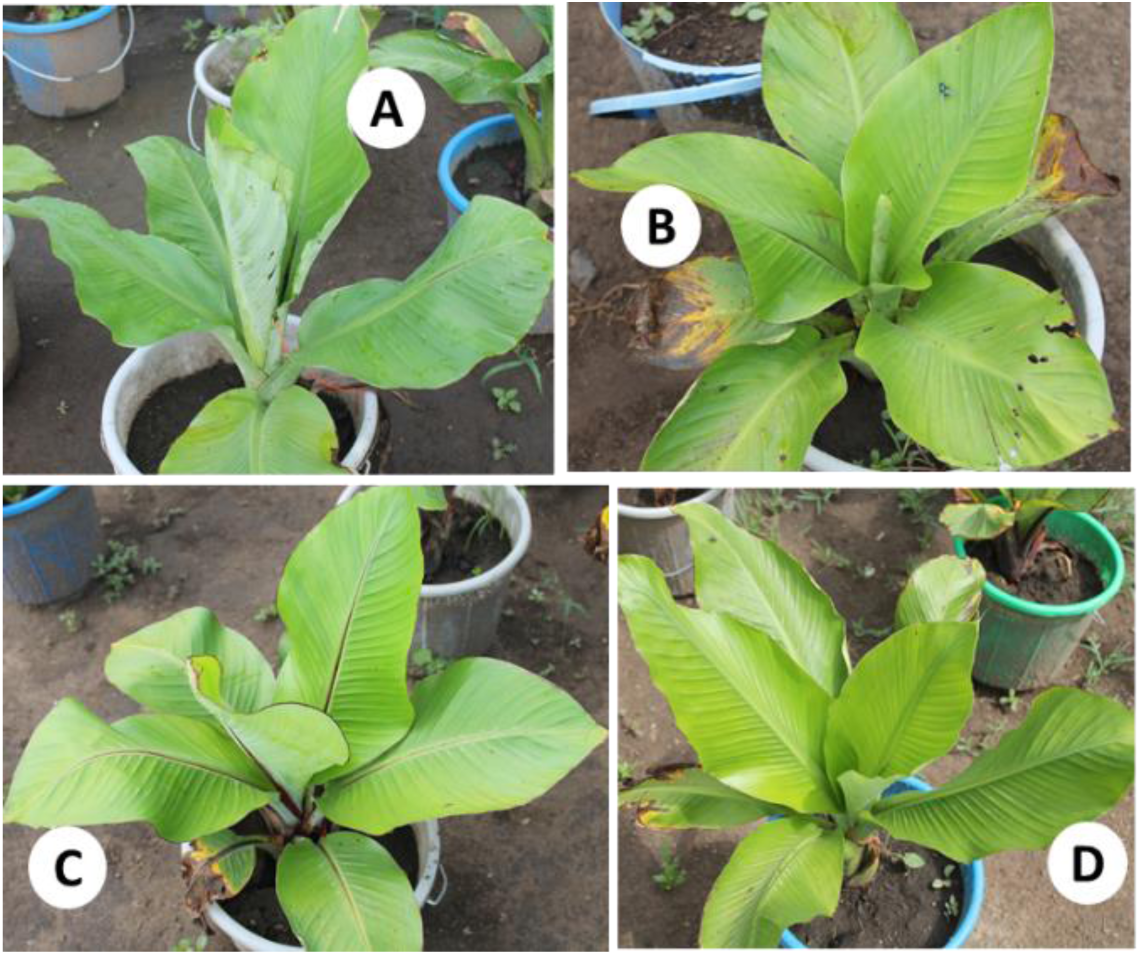
Xvm infection symptoms during the early stage of disease development on enset landraces. Labels are placed near to Xvm infected leaves. A) Curling of inoculated leaf, B) Twisting of inoculated leaf, C) Drooping of leaf apex of infected and other leaves and D) Folding around point of inoculation.

On severely infected enset landraces such as Dirbo and Arkia (susceptible control), the symptoms further developed into yellowing of leaves starting from leaf apex, and then gradual collapsing and clear wilting of the inoculated leaf and spreading the symptoms to other leaves. Eventually the whole leaves wilt, leading to their death and subsequent rotting of the whole plant (Figure 3A). However, on landraces that showed mild infection symptoms, for example Lemat and the resistant/tolerant control, Mazia, the inoculated leaf collapses and then dries (reminiscent of a classical hypersensitive response) or the symptoms extended to just a few adjacent leaves and the whole plant remains green afterward (Figure 4B). During the study period, a classical HR or HR-like symptoms were not observed in all of tested enset landraces.

**Figure 4.**
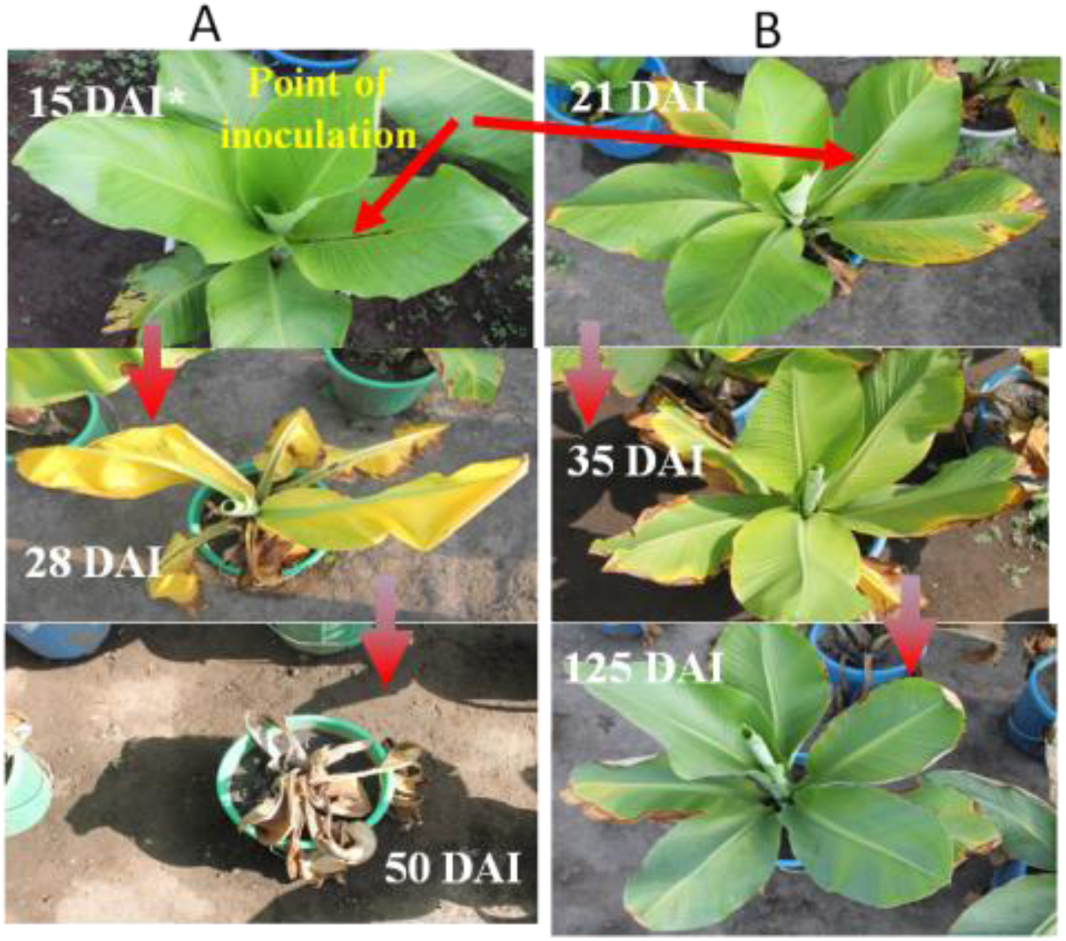
Comparison of Xvm disease symptoms on enset landraces. Development of Xvm disease sysmptoms on the susceptible enset landrace Dirbo (A) contrasted to the resistant/tolerant landrace Mazia (B).*DAI = Days after inoculation.

### 3.4. Area under disease progress stairs (AUDPS)

Plot of the area under disease progress stairs showed more or less distinct curves and disease build-up for the tested 20 enset landraces (Figure 5A). Analysis of AUDPS also showed a spectrum of significant differences (*P*<0.05) among landraces with landrace Arkia having the highest and Hae’la the lowest AUDPS value (Figure 5B). In addition, the analysis revealed that landraces could be grouped to fewer clusters based on their AUDPS values; those at lower tail, middle to upper tail, the last landrace Arkia that showed the peak AUDPS value.

**Figure 5.**
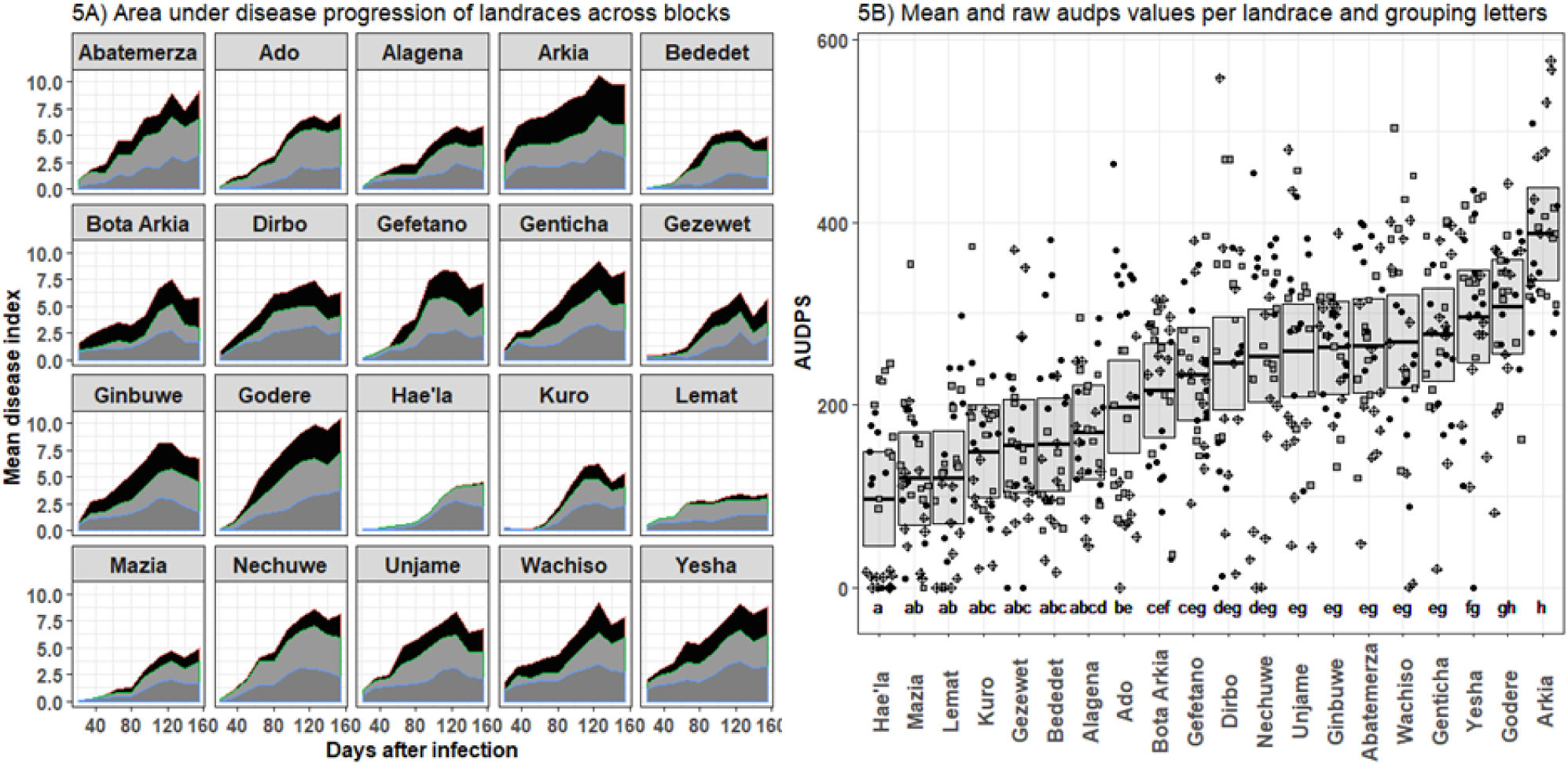
Area under the disease progression stairs (AUDPS) for 20 the landraces following controlled inoculation with Xvm. (A) Disease progression curve of tested enset landraces across blocks. The area under curve for each replicate in landrace represented by three shades of gray in each landrace. (B) Means (thick horizontal lines) and 95% confidence intervals (CIs) (shaded boxes) of the AUDPS values as estimated by a linear mixed effects model. Calculated areas for all individuals are plotted, with symbols indicating different replicates. The compact letter display above the landrace names indicates the results of multiple comparison tests using of means AUDPS using Tukey’s method at *P*<0.05. Enset landraces with similar letters are statistically not different from each other and vice versa.

### 3.5. Disease Index and apparent infection rate

Disease severity expressed as unit of DI illustrated the progress of disease development of Xvm infection in the 20 enset landraces. Apparent Infection Rate (AIR) calculates the rate of disease index development as a function of slope at each disease evaluation period (See Material and Methods). Analysis of variance of both the DI and AIR showed marked differences (P< 0.05) between landraces (Table 2).

**Table 2.** Analysis of variance for area under disease progress stairs, disease index and apparent infection rate of the twenty enset landraces infected with enset bacterial wilt disease.

|  | Degrees of freedom | Sum squares | Mean squares | F value |
| --- | --- | --- | --- | --- |
| Area Under Disease Progress Stairs (AUDPS) |  |  |  |  |
| Landrace | 19 | 3,176,321 | 167,175 | 19.29 *** |
| Disease Index (DI) |  |  |  |  |
| Landrace | 19 | 573 | 30 | 37.90 *** |
| Apparent Infection Rate (AIR) |  |  |  |  |
| Landrace | 19 | $1.04 \times 10^{-3}$ | $5.5 \times 10^{-4}$ | 7.18 *** |
| Residuals | 578 | 0.044 | $7.63 \times 10^{-5}$ | |

Mean DI values ranged from 0.65, 0.80, 0.81 in Hae’la, Mazia and Lemat respectively, corresponding to 16.25%, 20.00%, 20.25% severity. This contrasted with DI values of 1.98 (49.50%), 2.05 (51.25%), and 2.59 (64.75%) in Yesha, Godere and Arkia, respectively, (Table 3). As evidenced from the lowest AIR, disease development rate is the slowest in landrace Lemat, and fastest in Genticha, Abatemerza, Nechuw, Gefetano and Godere. Interestingly, AIR did not show a direct linear relationship with disease index suggesting that AIR varies among enset landraces irrespective of disease severity of landraces as measured by DI.

**Table 3.** Mean of disease index and apparent infection rates of Xvm on 20 enset landraces with their standard error values.

| Landraces | Disease index | cld<sup>1</sup> | Apparent infection rate<br>(AIR) | cld<sup>2</sup> |
| --- | --- | --- | --- | --- |
| Hae'la | 0.65 ± 0.06 | a | 1.3x10<sup>-2</sup> ± 0.002 | ac |
| Mazia | 0.80 ± 0.06 | ab | 1.4 x10<sup>-2</sup> ± 0.002 | acd |
| Lemat | 0.81 ± 0.05 | ab | 0.7 x10<sup>-2</sup> ± 0.002 | a |
| Kuro | 1.00 ± 0.06 | ac | 1.7 x10<sup>-2</sup> ± 0.002 | bce |
| Gezewet | 1.04 ± 0.07 | acd | 1.6 x10<sup>-2</sup> ± 0.002 | bce |
| Bededet | 1.05 ± 0.06 | acd | 1.5 x10<sup>-2</sup> ± 0.002 | ae |
| Alagena | 1.14 ± 0.06 | bcd | 1.5 x10<sup>-2</sup> ± 0.002 | ae |
| Ado | 1.32 ± 0.08 | ce | 1.9 x10<sup>-2</sup> ± 0.002 | bce |
| Bota Arkia | 1.44 ± 0.06 | def | 1.3 x10<sup>-2</sup> ± 0.002 | ab |
| Gefetano | 1.56 ± 0.08 | ef | 2.2 x10<sup>-2</sup> ± 0.002 | def |
| Dirbo | 1.64 ± 0.08 | efg | 1.4 x10<sup>-2</sup> ± 0.002 | ae |
| Nechuwe | 1.69 ± 0.09 | eh | 2.2 x10<sup>-2</sup> ± 0.002 | ef |
| Unjame | 1.73 ± 0.07 | fh | 1.6 x10<sup>-2</sup> ± 0.002 | bce |
| Ginbuwe | 1.76 ± 0.06 | fh | 1.6 x10<sup>-2</sup> ± 0.002 | bce |
| Abatemerza | 1.77 ± 0.08 | fh | 2.1 x10<sup>-2</sup> ± 0.002 | cef |
| Wachiso | 1.80 ± 0.08 | fh | 1.6 x10<sup>-2</sup> ± 0.002 | bce |
| Genticha | 1.85 ± 0.07 | fh | 2.0 x10<sup>-2</sup> ± 0.002 | bcef |
| Yesha | 1.98 ± 0.07 | gh | 1.8 x10<sup>-2</sup> ± 0.002 | bce |
| Godere | 2.05 ± 0.09 | h | 2.8 x10<sup>-2</sup> ± 0.002 | f |
| Arkia | 2.59 ± 0.06 | i | 1.5 x10<sup>-2</sup> ± 0.002 | ae |

cld<sup>1</sup> and cld<sup>2</sup> are compact letter display of the result of multiple comparison tests using of means of disease index and apparent infection rate using Tukey's method p<0.05. Enset landraces with similar letters are statistically not different from each other and vice versa.

### 3.6 Survival Analysis

A DI of ≥ 2.23 was used as a time-to-event cutoff point for survival analysis. This cut-off DI is the peak of infection level of the resistant/tolerant control enset landrace “Mazia”. The survival data generated with the survfit function of the survival package against landraces stratified by block, best fitted a Gaussian distribution that showed the minimum AIC and BIC values (Table 4) among Weibull, Logistic, Lognormal distributions. The linear fit of survival showed non-proportionality of hazard ratio and, alternatively, data fitted to a mixed effects Cox model for subsequent comparison of estimates (**supplementary R_script**).

**Table 4.**
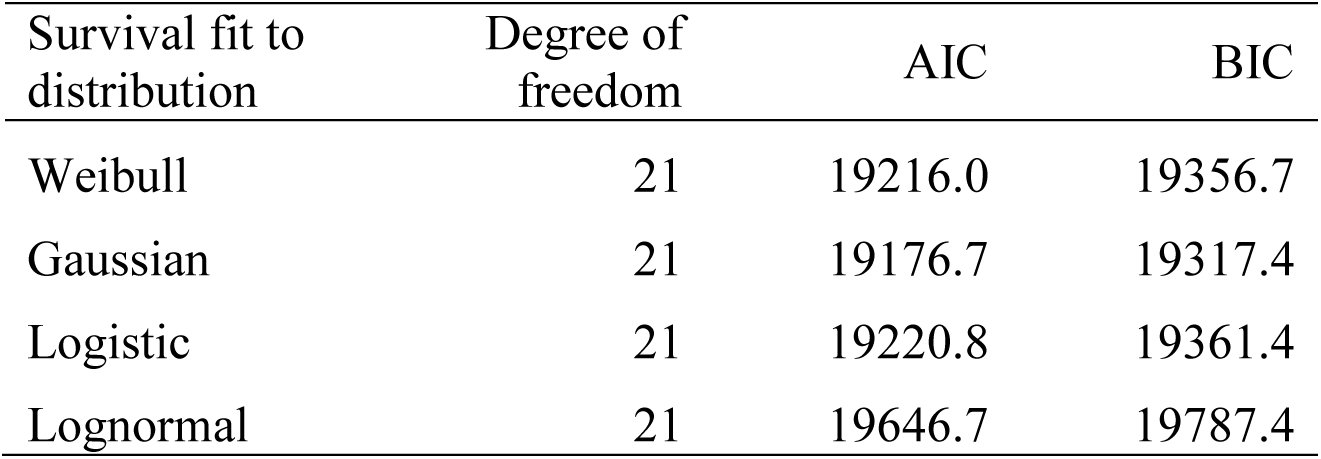
Summary of survival data fit to different distributions

| Survival fit to distribution | Degree of freedom | AIC | BIC |
| --- | --- | --- | --- |
| Weibull | 21 | 19216.0 | 19356.7 |
| Gaussian | 21 | 19176.7 | 19317.4 |
| Logistic | 21 | 19220.8 | 19361.4 |
| Lognormal | 21 | 19646.7 | 19787.4 |

Enset landraces displaying mild and severe infection to Xvm only showed significant estimates of hazard ratio (HR) and Gaussian distribution (Table 5). Lemat, Hae’la and Mazia had estimated hazard ratios of 0.230, 0.272 and 0.294, respectively, indicative of a mild infection to Xvm. This result indicates that for landraces Lemat, Hae’la and Mazia there is, respectively, an 87.0%, 86.8%, 80.6% decrease in risk of getting severely infected with Xvm higher than disease index (DI) value of ≥ 2.23 which was taken as a cutoff point. The susceptible control landrace Arkia included in this study showed an estimated hazard ration of 1.866 that suggests 86.6% higher risk of showing sever infection higher than DI value ≥ 2.23 after challenged with Xvm.

**Table 5.**
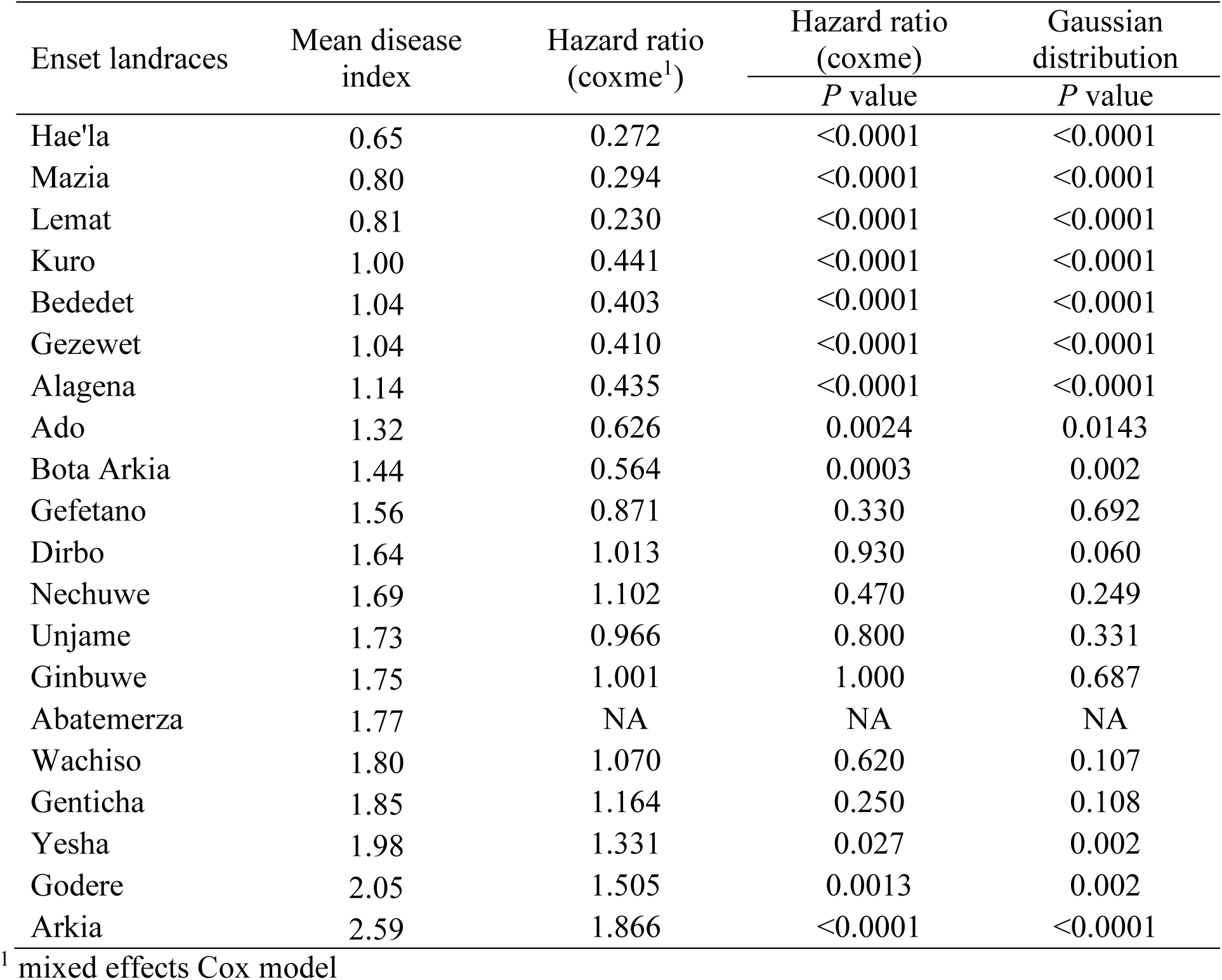
Result of Cox model and fit to Gaussian distribution of disease index of enset landraces

A survival fit to Gaussian distribution revealed that a higher proportion of plants survive from landraces that showed mild infection than those landraces that showed severe infection (see peak disease index) during the course of disease development (Figure 6). For example, right after 95 days post inoculation 50% of landrace Arkia was estimated to have experienced little probability of surviving whereas for Lemat, Hae’la and Mazia at this time point it was less than 7%.

**Figure 6.**
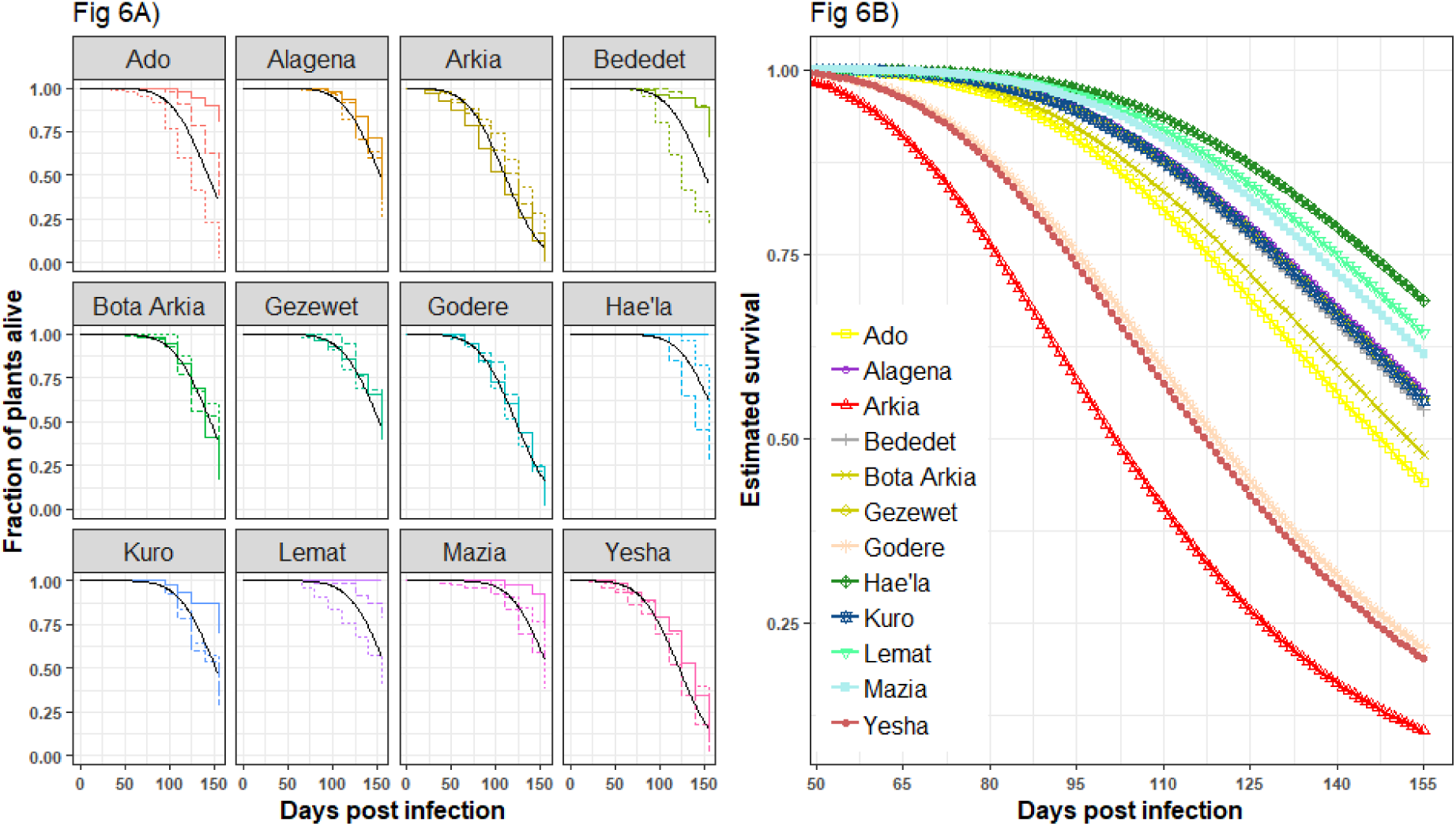
Kaplan-Meier estimates of survival and fits produced by survival regression analysis for disease index of the 12 enset landraces that fitted to Gaussian distribution. Landraces that showed non-significant values for hazard ratio and Gaussian distribution were omitted in this display. 6a) Kaplan-Meier survival estimates for individual landrace with different linetypes indicating experimental replicates. 6b) Combined display of Kaplan-Meier survival estimates for the 12 enset landraces. The event of interest was defined as a disease index of 2.23.

## 4. Discussion

The bacterial wilt disease caused by *Xanthomonas vasicola* pv. *musacearum* (formerly *X. campestris* pv. *musacearum*) is a major bottleneck to enset (*Ensete ventricosum*) cultivation in south and southwestern Ethiopia, where diverse enset landraces and wild types of the crop are found. Farmers traditionally adopted host plant resistance as a management option for this bacterial wilt disease by incorporating certain landraces perceived to be more resistant/tolerant to Xvm (Ashagari, 1985; Yemataw et al., 2017). Several studies have indicated the presence of enset landraces that show low level of susceptibility to Xvm (Ashagari, 1985; Archaido and Tessera, 1993; Handoro and Welde-Michael, 2007; Welde-Michael et al., 2008; Haile et al., 2014; Hunduma, 2015; Wolde *et al.*, 2016; Handoro and Said, 2016).

To date there has been no systematic comparison of enset landraces’ responses to Xvm infection in a common environment with a consistent inoculum. This detailed and, by nature of enset propagation, time consuming study undertook a detailed assessment of disease development on enset landraces previously reported for their low susceptibility to Xvm with the objective of gaining a definitive insight into Xvm disease phenotypes on enset landraces with the future objective of identifying suitable landraces to study the underlying molecular components of Xvm resistance. Characterisation of early infection symptoms through to the percentage of symptomatic leaves at each evaluation stage were considered. Various parameters such as area under disease progress stairs, disease index, apparent infection rate and survival rate were incorporated into detailed statistical analyses.

A limitation of this study was the unavailability of a standard stock of virulent inoculum. Given the constraints of available infrastructure in the geographic region of this study, the only practical possibility was to use freshly harvested inoculum from already infected plants. This approach was validated though preliminary experiments demonstrating that culturing the bacteria resulted in a significant attenuation of virulence (based on HR in tobacco and symptoms on Arkia); therefore, uncultured inoculum (i.e. bacterial suspension derived directly from bacterial ooze of infected plans) was used. Most previous enset-Xvm interaction studies have used Xvm isolates from disease hot-spot areas in southern Ethiopia in the District of Hagere Selam where highly virulent Xvm isolates were identified by Handoro and Welde-Michael (2007), based upon the observation that the tolerance/resistant landrace Mazia showed comparatively higher-level of infection to Xvm from Hager Selam districts than isolates from four other districts. However, comprehensive studies on classical and molecular epidemiology on diverse Xvm strains from enset growing areas in Ethiopia is required, and is currently underway, to identify virulent Xvm strains for future breeding efforts. Comparison of results between different studies has been hampered by the inevitable variability between batches of freshly collected inoculum. However, within our present study, we used a single uniform batch of inoculum, simultaneously inoculating all plants, thus eliminating such confounding batch effects.

The majority of vascular pathogens, including Xvm, manifest wilting symptom in their host after infection. Yet, even in well developed model systems, knowledge of the underlying infection mechanisms remain poorly resolved. Despite such challenges, it is important to understand the temporal spatial development of symptoms from initial visible phenotypes to the actual wilting and subsequent death. Furthermore, it is important to understand how different closely related genotypes respond to the same Xvm challenge. In this study we show that during early stages of Xvm infection on enset landraces, i.e. up to 15 days after inoculation, a range of transient symptoms were observed, including twisting and leaf curling, and drooping of the leaf blade and apex. Symptoms associated with wilting, clear folding or collapsing of leaf blade, severe drooping from leaf apex and wilting tended to appear from the third week following Xvm infection. Thus, for robustness of scoring, analyses were restricted to disease data gathered from the third week onwards.

This study revealed HR-like symptoms in a limited number of Xvm challenged enset landraces, reminiscent of that seen in resistance-gene mediated HR. This raises the possibility that disease classical resistance genes are deployed in restricting Xvm ingress. Involvement of HR cannot be excluded even in those landraces that do not show HR-like symptoms; evidence suggests that local resistant gene mediated reactions can also manifest as cellular changes contributing to wilting (Jakobek and Lindgren, 1993; Chasan, 1994; Pajerowska-Mukhtar and Dong, 2009). All 20 tested enset landraces showed at least partial symptoms of Xvm infection at different times of assessment while mock infected plants were asymptomatic during the course of evaluation. Destructive bacterial multiplication was not assessed at each disease evaluation stage, but rather the focus was to distinguish symptoms associated with disease and potential symptoms associated with an immune response. However, it is important to note that resistant plants also display a low level of infection symptoms as reported in several crops. Indeed, bacterial enumeration of most classical gene-for-gene interactions demonstrate a low level of accumulation of the pathogen until effective resistance sets in and this is probably reflected in the weak phenotypes recorded in these landraces. Previous reports which claim a lack of complete immunity to Xvm infection in enset due to the presence of some level of infection in tested landraces (Archaido and Tessera, 1993; Handoro and Welde-Michael, 2007; Welde-Michael et al., 2008; Haile et al., 2014; Hunduma, 2015; Wolde *et al.*, 2016; Handoro and Said, 2016) probably highlight the challenge in deploying effective resistance to a vascular pathogen compared to a more classical foliar infection.

Analysis of area under disease progress stairs and disease index discriminated those enset landraces with lower and higher level of infection, identifying elite landraces for further studies. Accordingly, landrace Hae’la, Mazia and Lemat showed markedly lower susceptibility to Xvm where as Yesha, Godere and Arkia were significantly more susceptible. Apparent infection rates indicated that disease development was slowest in Lemat and fastest in Yesha compared to the other landraces.

Complementing these results, survival analysis provided insight into the inherent infection risk. Inoculations leading to a cutoff disease index value of ≥ 2.23 corresponded to 55.75% disease incidence. The maximum disease value recorded was from the resistant control landrace Mazia and this was used as a reference in the survival analysis. The significant difference in proportional hazard model and Kaplan-Meier estimates of survival time demonstrated the variation in gross impact of Xvm in different enset landraces. Analysis of the hazard ratio suggested 2.4 to 3.9 fold changes in the resistance/susceptibility of landrace to disease caused by Xvm. Survival analysis suggested a higher proportion of landraces from Hae’la would be estimated to survive after Xvm infection followed by Lemat and Mazia whereas most plants in landrace Areki would succumb to Xvm infection, followed by landraces Yesha and Godere.

This study confirmed the lower level of susceptibility of previously reported enset landraces to Xvm infection after subjecting the landraces to a longer infection period, undertaking in parallel a detail account of symptomologies and rigorous analysis of disease data. It is important to mention that there is a great deal of evidence of sharing vernacular of enset landraces within and in neighboring enset growing districts in Ethiopia (Olango et al, 2015; Yemataw et al., 2014; 2016) that might not be phenotypical or genetically identical. For example, Olango et al. (2015) revealed by Simple Sequence Repeat based genetic diversity analysis that two enset landraces with identical vernaculars ‘Gena’ from Wolaita and Sidama zones in Southern Ethiopia were genetically distinct. Futhermore, genome-wide comparison of SNP data from the landrace Mazia sourced from Dawro and Wolaita zones of Southern Ethiopia revealed different SNP profiles (Yemataw *et al.*, 2018). Thus, care needs to be taken when identifying enset landraces for study as variation in the reaction of landraces with identical or similar vernaculars might not be only due to the variation in virulence of Xvm isolates or environmental factor but also due to of the presence possible homonym enset landraces.

Most of the landraces that showed higher tolerance/resistance to Xvm infection viz. Hae’la, Mazia, Lemat, Kuro, Bededet, and Gezewet originated from areas where landrace enset landrace diversity is richer with a high number of unique landraces present (Yemataw et al., 2014; 2016). These areas are the Dawro (landrace Mazia and Kurro), Kembata (landrace Hea’la), and Gurage (landrace Lemat, Bededet, and Gezewet) districts of southern Ethiopia. The recent disease pressure in these districts is comparatively higher than other enset growing areas in southern Ethiopia except for Kembata Tembaro district (SARI-McKnight CCRP, 2013; and personal observation by Sadik Muzemil). Furthermore, these areas also reside along the belts of the initial discovery of Xvm by Castellani (1939) in southern Ethiopia. Hence, we hypothesize that the co-existence of Xvm with these enset might have contributed to evolution of landraces with lower susceptibility to Xvm infection.

Enset landraces with lower susceptibility to Xvm identified in this study i.e landraces Hae’la, Lemat and Mazia, along with highly susceptible landraces provide the foundation for further testing in multi-stage and environment conditions in order to better understand the underpinning resistance/tolerance components. We note that it is important that not only the source of landraces, but also the pathogen inoculum need to be considered for future studies.

## Acknowledgement

The authors would like to thank McKnight Foundation Collaborative Research Program and the European Community Horizon 2020 grant Project ID 727624, “Microbial uptakes for sustainable management of major banana pests and diseases (MUSA) for financial and technical support. We thank Areka and Hawassa Agricultural Research Centers, Southern Agricultural Research Institute and Ethiopian Agricultural Research Institute for hosting and provision of germplasm as well as necessary services and facilities during field study. We are indebted to School of plant and horticultural science of Hawassa University for hosting the study, undertaken as part of SM’s MSc research work.

## Compliance with Ethical Standards’

The authors would like to undertake that all authors listed have contributed sufficiently to the project to be included as authors. To the best of our knowledge, no conflict of interest, financial or other, exists. We have included acknowledgements, conflicts of interest, and funding sources after the discussion.

